# Identification of a cis-acting determinant limiting expression of sphingomyelinase gene *sph2* in *Leptospira interrogans* with a *gfp* reporter plasmid

**DOI:** 10.1101/400259

**Authors:** James Matsunaga, David A. Haake

## Abstract

Many Pomona strains of the spirochete *Leptospira interrogans* express the osmotically-inducible sphingomyelinase gene, *sph2*, at much higher levels than non-Pomona strains. We developed a new green fluorescent protein (GFP) reporter plasmid to examine *sph2* gene expression determinants. The vector allows fusion of the test promoter to the ribosome-binding site and coding region of *gfp*. We fused the *sph2* promoters from the serovar Lai strain 56601 and from the serovar Pomona strain LC82-25 to *gfp* to examine the molecular determinants of differential *sph2* expression between the two strains. Similar to what is observed with the native *sph2* genes, introduction of the plasmids into the Lai 56601 strain resulted in near background levels of *gfp* expression from the Lai *sph2* promoter, while expression from the Pomona *sph2* promoter was high. Expression from both fusions increased at physiologic levels of osmolarity achieved by addition of sodium chloride to the culture medium. We examined the role of a 17 bp upstream element found in all *L. interrogans* strains expressing low basal levels of *sph2* and missing from Pomona strains that express *sph2* at high levels. When the 17 bp sequence present upstream of the Lai *sph2* promoter was deleted or scrambled, fusion expression increased substantially. Conversely, insertion of the 17 bp sequence upstream of the Pomona *sph2* promoter diminished fusion expression. In contrast, removal of an insertion sequence-like element that is found only in the Pomona *sph2* upstream sequence had no effect on expression from the Pomona *sph2* fusion in the Lai strain. These findings demonstrate the utility of the *gfp* reporter plasmid in analyzing gene expression in *L. interrogans*.

**IMPORTANCE**

Genetic tools are needed to examine gene expression in the pathogen *Leptospira interrogans*. We developed a reporter plasmid that replicates in *L. interrogans* with green fluorescent protein (GFP) as the read out of promoter activity. We demonstrated an application of the new reporter plasmid by identifying an upstream element responsible for the poor basal expression of the *sph2* sphingomyelinase gene in a serovar Lai strain of *L. interrogans*. This new tool is useful for discovery of the molecular determinants of *L. interrogans* gene expression.

## INTRODUCTION

Most pathogenic members of the genus *Leptospira* have a bimodal lifestyle. In one phase, leptospires live in moist soil and freshwater bodies located throughout the world (1). At other times, they invade vertebrate hosts to permanently colonize the renal tubules, resulting in long-term shedding into the environment by these carriers via their urine (2). Adaptation to these diverse conditions requires regulation of large numbers of specialized genes, consistent with whole-genome transcriptional array and sequencing studies of *L. interrogans* incubated under conditions simulating those encountered during its life cycle (3-8). Additionally, different serovars of *Leptospira* are generally associated with specific hosts (9), which may reflect adaptation by differential expression of the same genes in different serovars of *Leptospira*.

One example of a gene expressed at dissimilar levels in different serovars is *sph2*, which encodes an enzyme responsible for most, if not all, of the sphingomyelinase activity of *L. interrogans* (10). *L. interrogans* expresses *sph2* in dialysis membrane chambers implanted into the peritoneal cavity of rats. Additionally, antisera raised against Sph2 detects antigen in kidney sections from acutely-infected hamsters (11) and in urine obtained from leptospirosis patients (12). Sph2 may contribute to the pathogenesis of leptospirosis by adhering to fibronectin, inducing apoptosis of host cells, and triggering the production of proinflammatory cytokines (13-16). Most members of serovar Pomona examined to date produce large amounts of *sph2* transcript and Sph2 protein, whereas a strain of serovar Lai produces lower amounts of transcript and undetectable levels of Sph2 during growth in conventional culture medium (10, 11,). Strains of serovars Copenhageni and Manilae also produce negligible amounts of Sph2 (10).

The genetic basis for the increased expression of *sph2* in Pomona strains is unknown. A molecular epidemiology study with PCR primers encompassing an insertion sequence-like element located upstream of *sph2* found that the element is widespread among Pomona subtype kennewicki strains (17). The IS-like element is missing from the *sph2* upstream sequences of the four *L. interrogans* strains shown to produce low basal levels of Sph2 (10). For these reasons, we hypothesized that this element is responsible for the high levels of Sph2 produced in the Pomona strains (10). However, it should be noted that this hypothesis was based on observations of a single Pomona strain (10), and Sph2 was not detected in another Pomona strain that has the IS-like element upstream of the coding region (13). Whether other Pomona strains that produce high levels of Sph2 harbor the IS-like element is unknown (11).

Several reporter constructs have been developed for studying the expression of *L. interrogans* genes such as *sph2*. Osmotic induction of the *ligA/ligB* and *sph2 L. interrogans* promoters was demonstrated by genetic fusions to *gfp* in the nonpathogen *L. biflexa* (18). The *kdp* promoter was fused to the 5’ end of the *L. biflexa* β-galactosidase gene *bgaL* and integrated by homologous recombination into the endogenous *bgaL* gene to create a model system for demonstrating positive regulation by KdpE of *L. interrogans* (19). The β-galactosidase gene from *Geobacillus stearothermophilus*, *bgaB*, was fused downstream of the *lig* promoter and 5’ untranslated region to examine expression in *E. coli*. Mutational analysis of the 5’ UTR showed that secondary structure sequesters the ribosome-binding site (20). In each of these cases, regulation of pathogenic gene expression was examined in a surrogate host because a plasmid able to replicate in *L. interrogans* was not available. Promoter fusions to *gfp* or luciferase genes have been introduced into *L interrogans* by transposition (21-23). However, mutational analysis of promoters fused to the reporter is not possible with the transposon-based system; random insertion of the transposon carrying the fusion complicates interpretation of expression levels because the location of the transposon in the chromosome may influence fusion expression. To overcome this limitation, we took advantage of the fact that several replicative plasmids for *L. interrogans* have recently been developed (24, 25,). For example, pMaORI was constructed by cloning the *rep* and partition loci from phage-like sequences integrated in the *L. mayottensis* genome (24). pMaORI has been shown to be stable in the absence of antibiotic selection in seven strains of *L. interrogans*, the nonpathogen *L. biflexa*, and the intermediate species *L. fainei* and *L. licerasiae*. Here, we adapted the pMaORI plasmid to create a new *gfp* reporter vector that can be used to examine gene expression in *L. interrogans*. We used the *gfp* reporter construct to identify a sequence responsible for the low basal levels of *sph2* expression in a Lai strain of *L. interrogans* as compared to that in a Pomona strain that produces large amounts of Sph2.

## MATERIALS AND METHODS

### Bacterial strains

*Leptospira interrogans* serovar Lai strain 56601 (26), *L. interrogans* serovar Copenhageni strain Fiocruz L1-130 (27), and *L. interrogans* serovar Pomona subtype kennewicki strains LC82-25, RM211, and P10637-46 (National Animal Disease Center, Ames, IA) were grown in EMJH (Millipore, sold as “Probumin Vaccine Grade Solution,” cat # 840665, lot 103) at 30°C with 40 µg/ml spectinomycin added for plasmid selection. *E. coli* strains DH5α, π1, and β2163 (28) were incubated in LB with the appropriate antibiotic added to select for plasmids. LB media for *E. coli* π1 and β2163 was supplemented with 0.3 mM thymidine and 0.3 mM diaminopilemic acid, respectively (28).

### Western blots

Immunoblot analysis of *L. interrogans* lysates was performed as described (20). Blots were probed with 1:1,000 dilution of Sph2 antiserum and 1:5,000 dilution of LipL41 antiserum (29, 30,).

### Plasmid construction

Plasmids used in the study are compiled in Table 1. Plasmid DNA was extracted from 4 ml of overnight *E. coli* cultures with the Qiagen QIAprep Spin Miniprep Kit and eluted with 50 µl of 10 mM Tris-HCl, 0.1 mM EDTA, pH 8.5 (Valencia, CA). Genomic DNA was purified from 5 ml of saturated *L. interrogans* cultures with the Promega Wizard Genomic DNA Purification Kit and resuspended in 50 µl of 1 mM Tris-HCl, 0.1 mM EDTA, pH 7.4. Restriction enzymes and Quick Ligase were supplied by New England Biolabs (Ipswich, MA). Synthetic oligonucleotides were obtained from Invitrogen, and the nucleotide sequences are provided in Table 2. PCR reactions were done with *Pfu* DNA polymerase following the manufacturer instructions (Thermo Fisher).

**Table 1.**
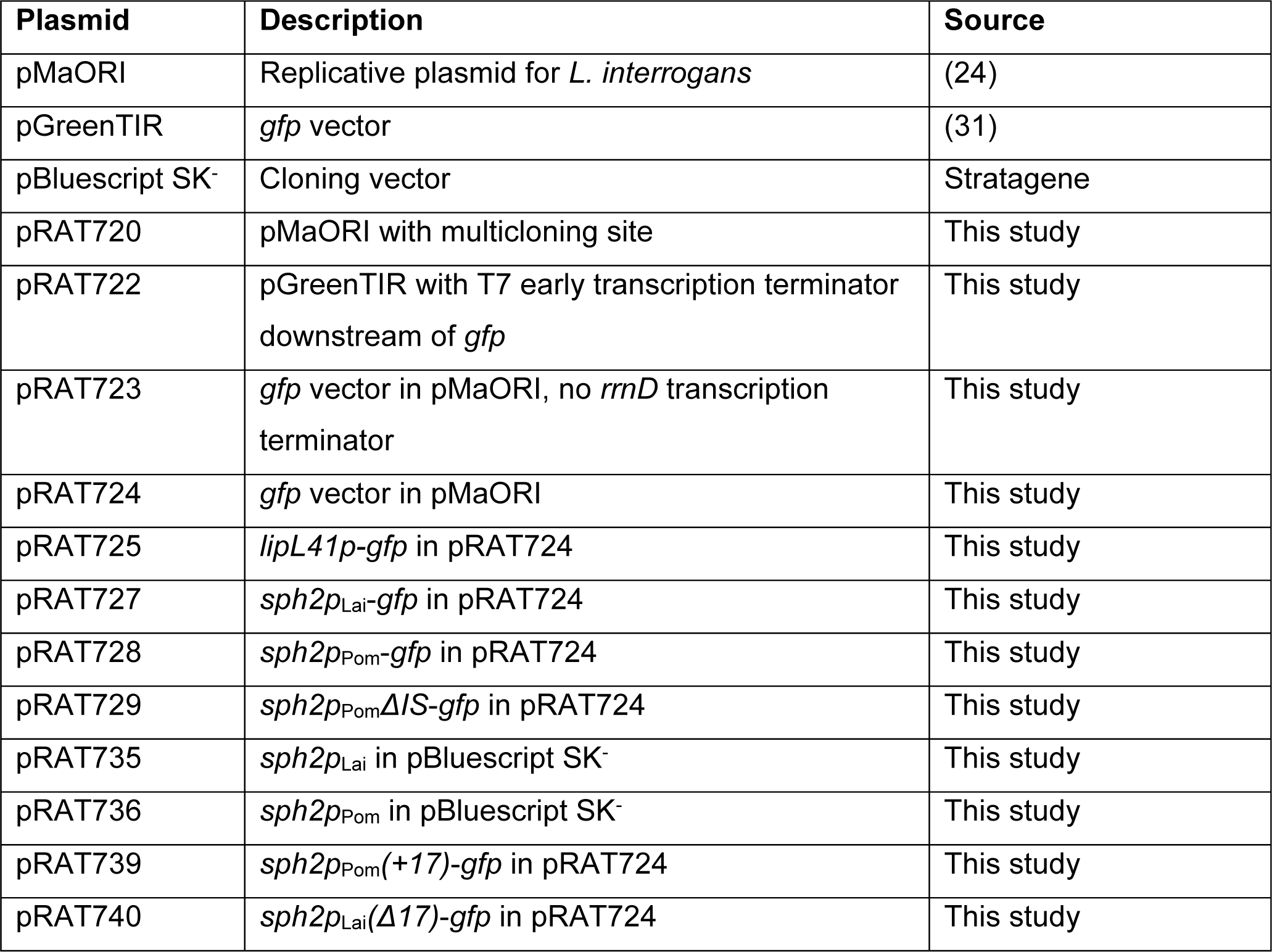
Plasmids used in this study

**Table 2.**
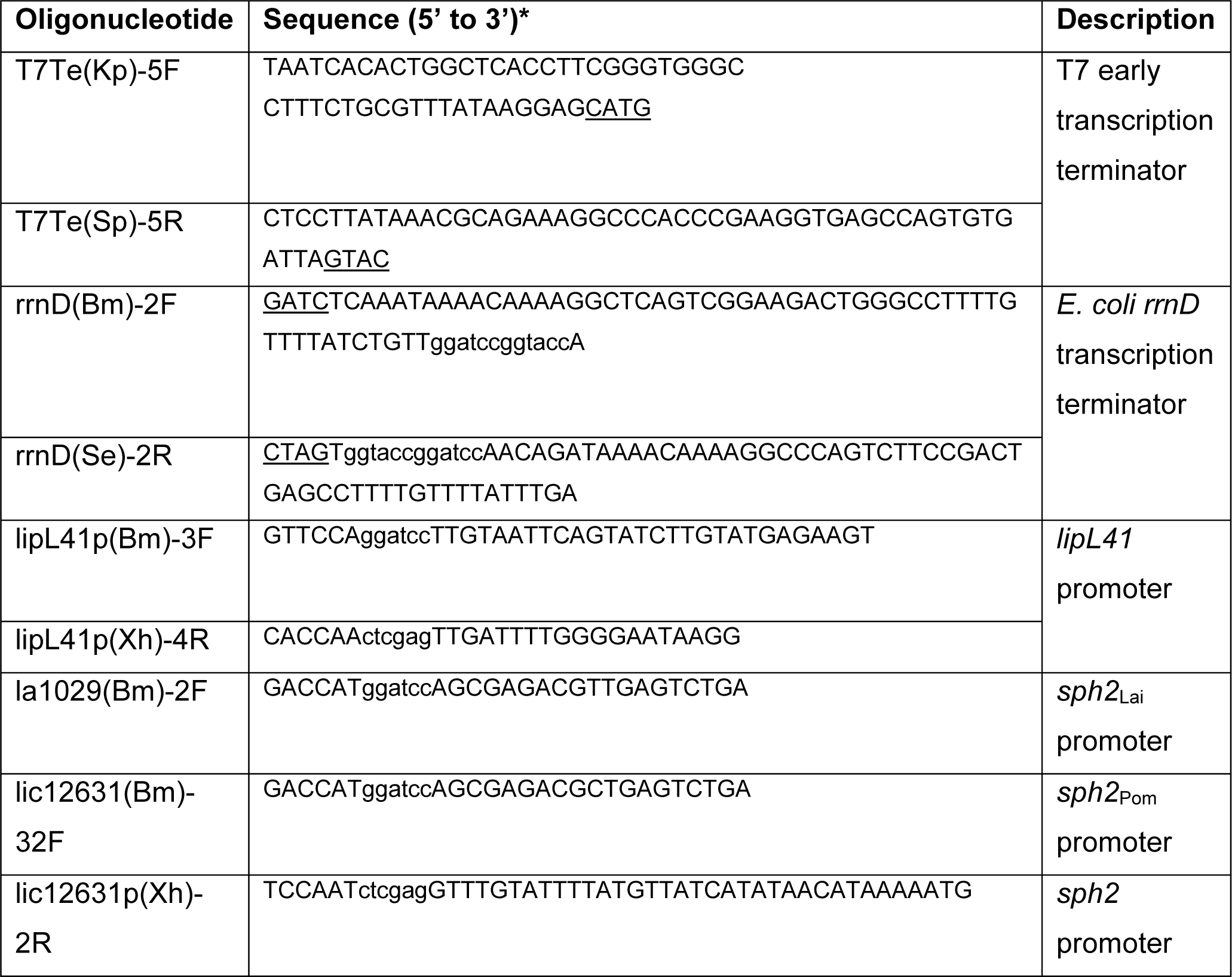
Synthetic oligonucleotides used for plasmid constructions

The *gfp* reporter plasmid was generated as follows (Fig. 1). A double-stranded oligonucleotide harboring the T7 early transcription terminator with 5’ *Kpn*I and 3’ *Sph*I single-stranded overhangs (Table 2) was inserted downstream of *gfp* in pGreenTIR (31) to create pRAT722. Additionally, a multicloning site was introduced into the plasmid pMaORI (24) by replacing its small *Bam*HI-*Sal*I fragment with the double-stranded DNA 5’-GATCCGGTACCACTAGTCTCGAGCCCGGGGCATGCG-3’/5’-TCGACGCATGCCCCGGGCTCGAGACTAGTGGTACCG-3’ (*Bam*HI and *Sal*I overhangs underlined), resulting in pRAT720. The *gfp* ribosome-binding site and coding region along with the 3’ T7 early terminator was excised from pRAT722 and inserted into *Xma*I-*Sph*I-cut pRAT720 to generate pRAT723. Finally, a double-stranded oligonucleotide carrying the *E. coli rrnD* transcription terminator with 5’ *Bam*HI and 3’ *Spe*I sticky ends and *Bam*HI and *Kpn*I sites 3’ of the terminator was inserted upstream of *gfp* sequences in pRAT723 to generate pRAT724.

**FIG 1.**
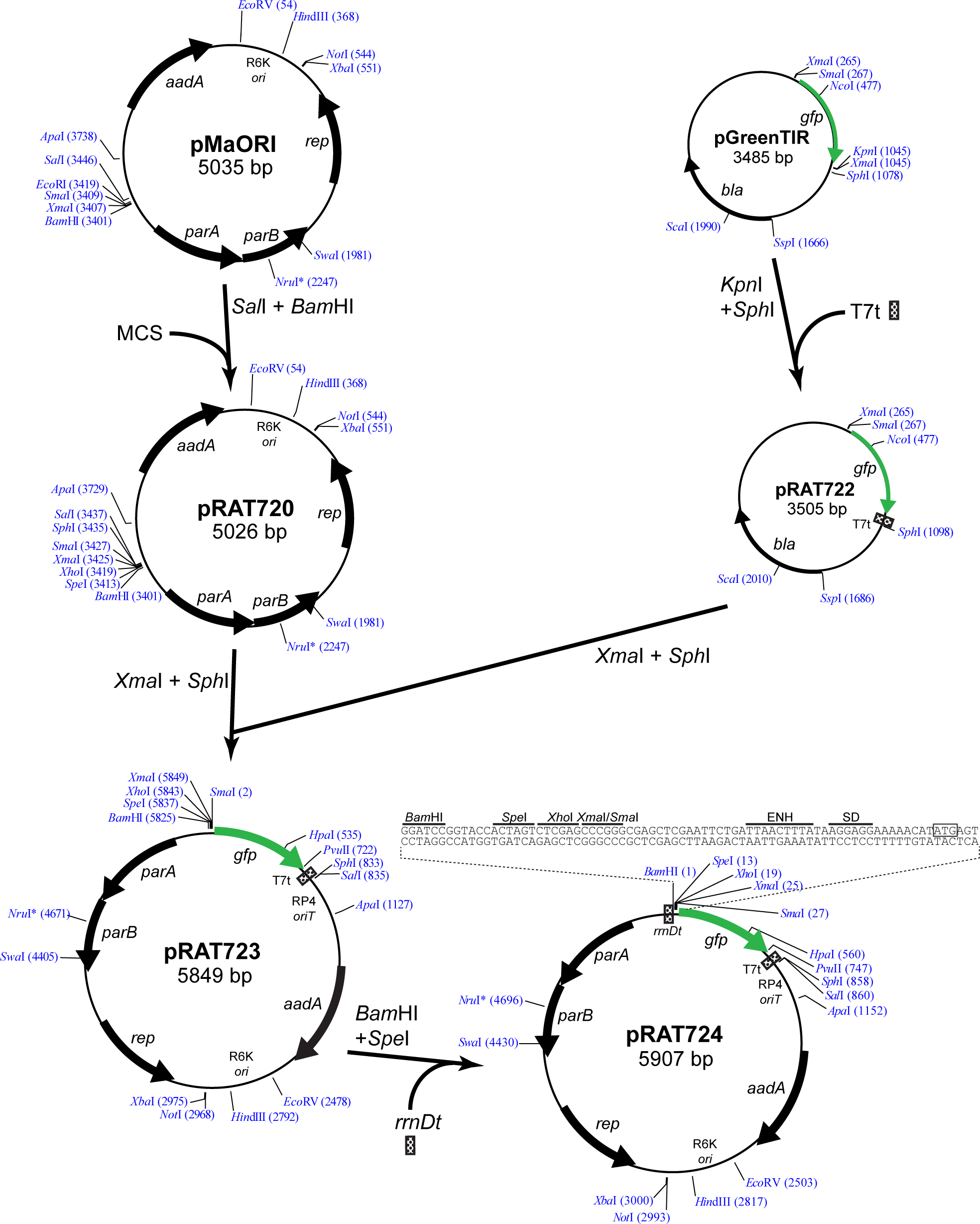
Construction of *L. interrogans gfp* reporter plasmid. Thick arrows denote important open reading frames. Restriction sites shown are unique except for the *Xma*I sites of pGreenTIR. Numbers next to restriction enzyme names refer to the nucleotide position following the cleavage site. Transcription terminators and a multicloning site were inserted as double-stranded oligonucleotides with overhangs compatible with the ends of the restriction digested parent plasmid DNA. *aadA*, spectinomycin resistance; *parA* and *parB*, partition locus; *rep*, replication initiator; ENH, T7 gene 10 translational enhancer; SD, Shine-Dalgarno sequence; T7t, early T7 transcription terminator; *rrnDt*, *E. coli rrnD* transcription terminator; RP4 *oriT*, RP4 origin of transfer; R6K *ori*, R6K origin of replication; MCS, multicloning site; *, restriction site subject to *dam* methylation.

To clone the *lipL41* and *sph2* promoters, PCR primers were designed with *Bam*HI and *Xho*I restriction sites near the 5’ ends of the upstream and downstream primers, respectively (Table 2). The *lipL41* promoter was PCR amplified using *L. interrogans* strain Fiocruz L1-130 genomic DNA as template. The *sph2* promoter was amplified using *L. interrogans* strain 56601 (sv. Lai) and strain LC82-25 (sv. Pomona) genomic DNA as template. PCR products were digested with *Bam*HI and *Xho*I and inserted into the corresponding site in pRAT724 to create the promoter fusions to *gfp*.

### Mutagenesis

The template for oligonucleotide-directed mutagenesis of the sequence upstream of the *sph2* promoter was generated by transferring the *Bam*HI-*Xho*I fragment containing the promoter sequence from the wild-type *sph2-gfp* fusion plasmids into pBluescript SK^-^ (Stratagene) to generate pRAT735 and pRAT736 (Table 1), which harbor the *sph2* promoters from the Lai 56601 and Pomona LC82-25 strain, respectively. The desired mutations upstream of the *sph2* promoter were generated as described (20, 32,). In brief, overlapping oligonucleotides with the desired changes directed towards each strand (Table 3) were used to PCR amplify the plasmid with *Pfu* DNA polymerase. The template DNA was depleted by digestion with *Dpn*I, and the amplicon was transformed into *E. coli* DH5α. The transformation culture was plated on LB plates containing 100 µg/ml ampicillin. Plasmid DNA was purified from overnight cultures of colonies, and the presence of the desired mutations was confirmed by Sanger sequencing with the T7 and T3 sequencing primers (Laragen, Culver City, CA).

**Table 3.**
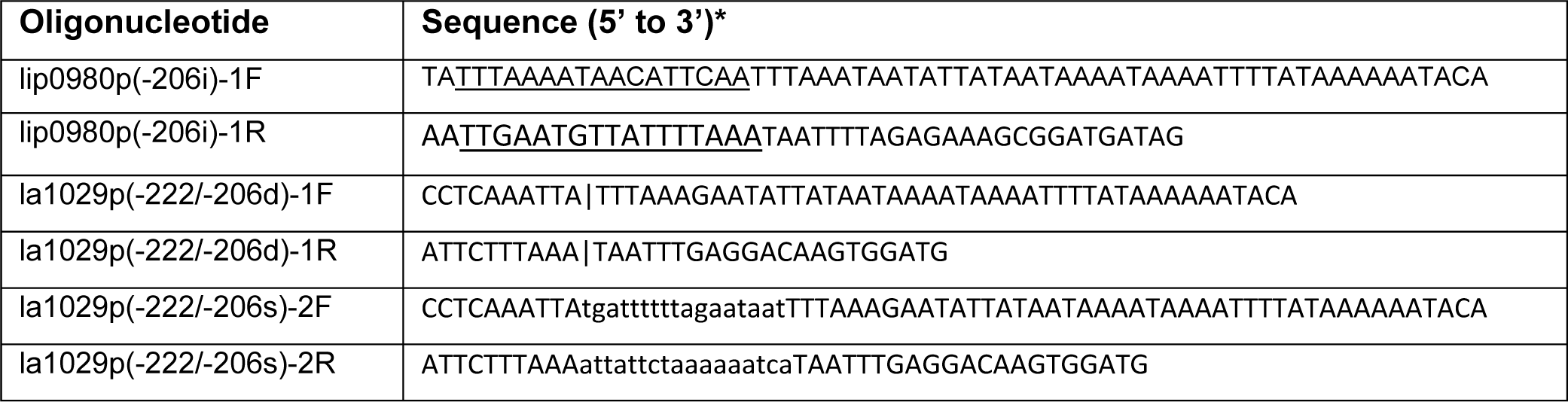
Oligonucleotides used for site-directed mutagenesis

### Conjugation

The *gfp* fusion plasmids were transferred into *L. interrogans* by conjugation from a donor *E. coli* strain as described (Slamti 2012). Briefly, *E. coli* β2163 was transformed with the *gfp* fusion plasmids and grown overnight in LB with 0.3 mM DAP and 40 µg/ml spectinomycin at 37°. The next day, 40 µl of the overnight culture was transferred to 4 ml of LB with DAP and grown to late exponential phase. *L. interrogans* was grown to a density of ≈10^8^/ml. 5 ml of *L. interrogans* and 0.5 ml *E. coli* were mixed and filtered with a 0.1 µm filter (Fisher). The filter was placed face up on an EMJH plate containing 0.3 mM DAP and incubated overnight at 30°C. The bacteria were washed off the filter with 1 ml of EMJH and plated onto three EMJH plates containing 40 µg/ml spectinomycin.

### 5’ end mapping of RNA

A culture of *L. interrogans* Fiocruz L1-130 was started at 2 x 10^7^/ml in 25 ml EMJH medium supplemented with 80 mM NaCl and incubated for 5 days to a density of 6 x 10^8^/ml. Additionally, a culture of L. interrogans LC82-25 was initiated in 25 ml EMJH at 1 x 10^7^/ml and incubated for 3 days to a density of 5 x 10^8^/ml. RNA was extracted from *L. interrogans* with Trizol (Invitrogen), and DNA was removed with Turbo DNase (Ambion). The 5’ end of the *sph2* transcript was determined with the Roche 5’/3’ RACE Kit, second generation (Roche) using oligonucleotide sph2-8R (5’-CGTTTTGCTCTTTCATCGTGTC-3’) to prime the reverse transcription reaction and sph2-9R (5’-GCGGAGCTGCCATTTTCTG-3’) to PCR amplify the dA-tailed *sph2* cDNA. The amplicon was sequenced with the sph2-9R primer to map the 5’ end.

### GFP assay

*L. interrogans* transconjugants carrying *gfp* fusion plasmids were grown in EMJH with 40 µg/ml spectinomycin to a density of ≈1-5 x 10^8^/ml, and the optical density of the cultures was measured at 420 nm (OD_420_) with an Ultrospec 2000 spectrophotometer (Pharmacia Biotech). 800 µl of each culture was collected by centrifugation for 5 min at 9,000 x g in an Eppendorf 5424 microcentrifuge. Cells were resuspended in a volume of PBS (OD_420_ x 4 ml) to obtain an OD_420_ reading of 0.4. 100 µl of each cell suspension was transferred to a dark-walled clear bottom 96-well Costar plate (Thermo Fisher), and the fluorescence emitted at 528 nm (excitation 485 nm) was measured with a Synergy2 Multi-Mode Microplate Reader (BioTek; Winooski, Vermont, USA). Because fluorescence readings of wild-type *L. interrogans* were similar to those of PBS, PBS was used as a blank for background subtraction.Fluorescence values from PBS were subtracted from readings of *L. interrogans* carrying the *gfp* fusions plasmids.

### Multisequence alignment

The *sph2* promoter sequences from *L. interrogans* strains Fiocruz L1-130, L495, 56601, and LC82-25 were aligned with Kalign (33).

### Statistics

Experiments were conducted with cultures initiated from three colonies (n = 3), unless stated otherwise. Statistical analysis was conducted with R, version 3.4.2, “Short Summer” (34).

## RESULTS

Previously, LC82-25 had been the only strain of serovar Pomona strain with an IS-like element upstream of *sph2* shown to produce high levels of Sph2 (10). For this reason, we tested two additional Pomona strains with the IS-like element for Sph2 production by Western blot analysis (Fig. 2). Both the RM211 and P10637-46 strains were isolated from pigs. The immunoblot shows that both strains, like the LC82-25 strain, produced high levels of Sph2 when incubated in EMJH. The levels of Sph2 were much higher than that produced by the *L. interrogans* serovar Manilae strain L495 strain grown in EMJH with sodium chloride added to achieve physiologic osmolarity.

**FIG 2.**
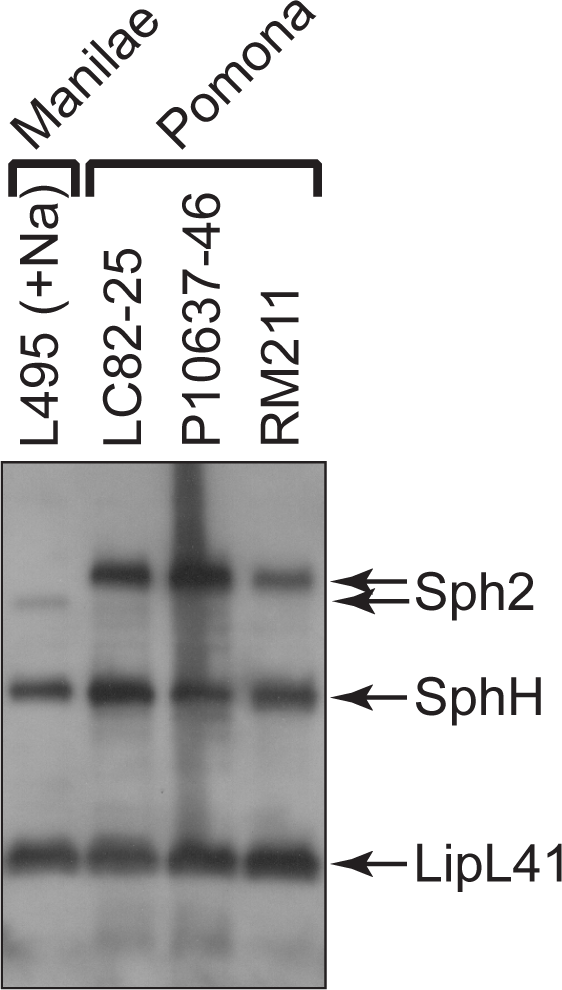
Western blot analysis of Sph2 production from three strains of *L. interrogans* serovar Pomona. Western blots from three *L. interrogans* Pomona strains grown in EMJH were probed with Sph2 and LipL41 antisera. The Sph2 antiserum cross-reacts with SphH. *L. interrogans* serovar Manilae strain L495 was grown in EMJH supplemented with 120 mM sodium chloride, and its lysate was included for comparison of Sph2 levels. Sph2 produced by the L495 strain (marked by *) has a slightly lower molecular weight than Sph2 produced by Pomona strains.

To determine whether transcription from within or adjacent to the IS-like element accounted for high basal expression of *sph2* in the Pomona LC82-25 strain, we mapped the major 5’ end of the *sph2* transcript by 5’ rapid amplification of cDNA ends (RACE) (Fig. 3A). The trace shows a single 5’ end located 28 nucleotides from the *sph2* start codon in both *L. interrogans* strain LC82-25 grown in EMJH and strain Fiocruz L1-130 grown at physiologic osmolarity (Fig. 3A). The mixture of the four nucleotides positioned next to the 5’ end likely stems from the terminal transferase activity of some reverse transcriptases (35). The 5’ end of the *sph2* transcript was found to be located downstream of a putative -10 and -35 *E. coli*-like promoter sequences (Fig. 3B).

**FIG 3.**
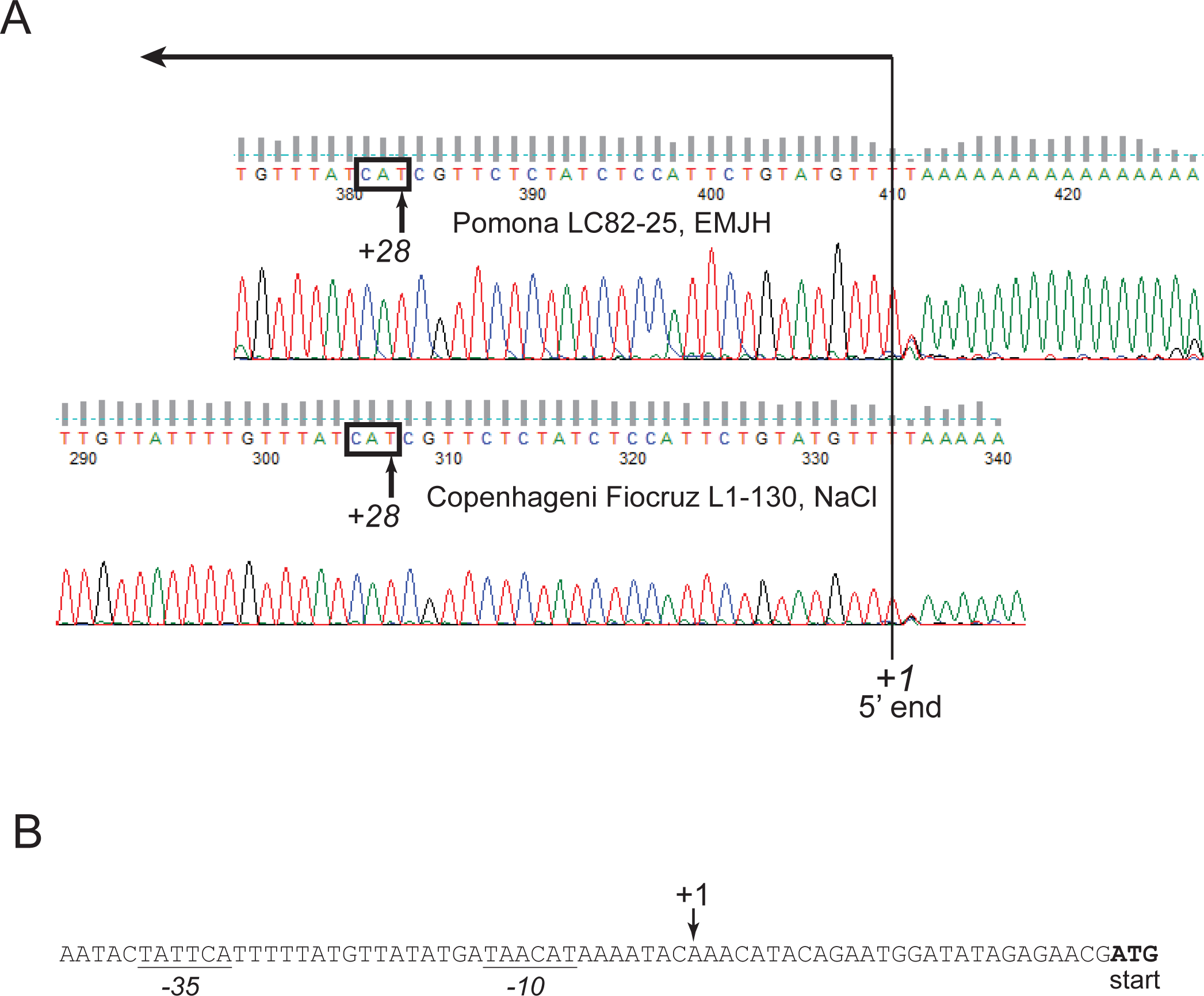
Mapping the major 5’ end of *sph2* by 5’ RACE. A. The 5’ ends of the *sph2* transcripts from *L. interrogans* LC82-25 grown in EMJH (top) and *L. interrogans* Fiocruz L1-130 grown in EMJH with 80 mM sodium chloride (bottom) were mapped by 5’ RACE. Electropherograms from sequencing reactions of the amplicons are shown. The peaks representing the major 5’ end of the *sph2* transcripts are bisected by a vertical line. The sequence complementary to the start codon is boxed. The direction of transcription is indicated by the long arrow. B. The nucleotide sequence around the major 5’ end of *sph2* is shown. The start codon is shown in bold, and the predicted -10 and -35 promoter elements are underlined.

We next constructed a *gfp* reporter plasmid to examine *sph2* gene expression in *L. interrogans*. We selected the *gfp* allele demonstrated previously to produce functional GFP in *L. biflexa* when fused to leptospiral promoters (18, 21,). The *gfp* allele includes the S65T “red-shift” mutation and the F64L mutation that increases protein solubility (31) and includes a Shine-Dalgarno sequence complementary to the 3’ end of the *L. interrogans* 16S rRNA. To construct the reporter plasmid, we cloned the *gfp* gene, along with its translation initiation region, into the plasmid pMaORI, an RP4-based mobilizable plasmid that replicates in *L. interrogans* (24). Five unique restriction sites are available for cloning promoters upstream of *gfp* (Fig. 1). A T7 transcription terminator was inserted downstream of *gfp* to create pRAT723 (Fig. 1).

To minimize transcription of *gfp* initiating from cryptic vector promoters upstream of *gfp*, an *rrnD* transcription terminator was placed upstream of the multicloning site. *E. coli* transformed with pRAT723, which lacked the transcription terminator, or pRAT724, which possessed the terminator, was collected by centrifugation of liquid cultures to examine the color of the cell pellet. To quantify the terminator’s activity in *L. interrogans*, pRAT723 and pRAT724 were transformed into an *E. coli* strain harboring the RP4 conjugation machinery and transferred by conjugation into the serovar Lai 56601 strain of *L. interrogans*. Cultures were grown from four colonies of each transconjugant and were processed for fluorescence measurements. The *rrnD* terminator reduced GFP production by ∼50% in both *L. interrogans* strains (Fig. 4). GFP levels in the absence of the *rrnD* terminator were similar to the levels produced from the *L. interrogans lipL41* promoter (Fig. 4).

**FIG 4.**
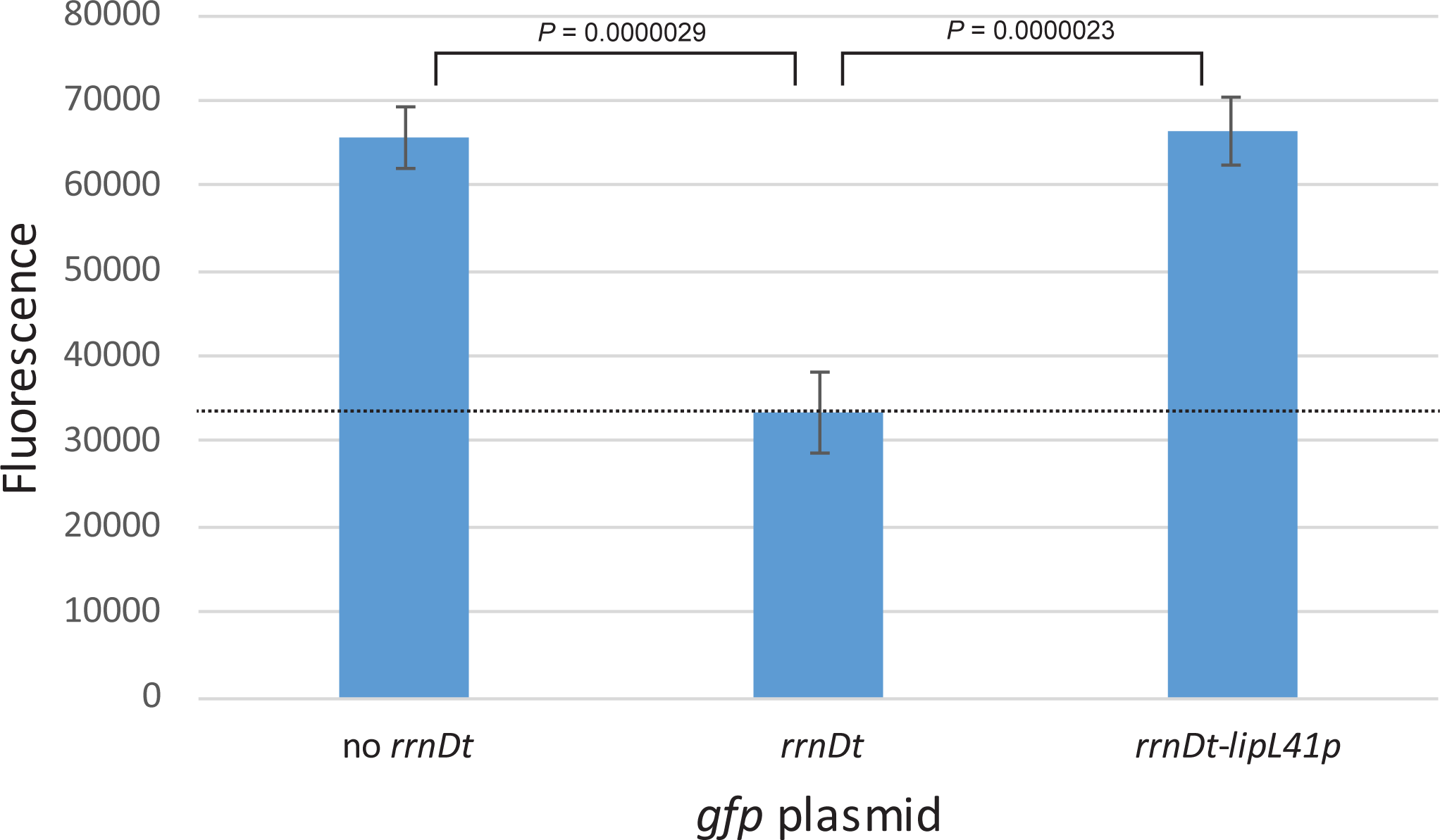
Effect of a transcriptional terminator on background *gfp* expression. Fluorescence measurements of *L. interrogans* containing plasmids with and without the *E. coli rrnD* transcription terminator upstream of *gfp* are shown. GFP produced from a *lipL41p-gfp* fusion with an upstream *rrnD* transcriptional terminator is shown for comparison. The dashed line indicates the mean background reading obtained with *L. interrogans* carrying the empty *gfp* fusion vector pRAT724. Means ± SD are plotted (n = 4). *P*-values were calculated by one-way ANOVA with Tukey HSD post hoc test.

The *sph2* transcript synthesized by the Lai 56601 strain is likely to have the same 5’ end as the *sph2* transcript produced by the Fiocruz L1-130 strain as the sequence upstream of the coding region in the two strains is nearly identical. Based on the 5’ RACE result, we cloned the promoter region of *sph2* from the Lai 56601 and Pomona LC82-25 strains upstream of *gfp* in pRAT724. The *sph2* sequence from the Lai strain comprised nucleotides -298 through +3 relative to the major 5’ end of the transcript. The *sph2* promoter fragment cloned from the Pomona strain had identical 5’ and 3’ ends as the Lai fragment and included the IS-like element. The *lipL41* promoter was also cloned into pRAT724. The resulting plasmids carrying the *sph2p-gfp* and *lipL41p-gfp* fusions, along with pRAT724, were introduced into the Lai and Pomona strains by conjugation. The four plasmids were also introduced into the nonpathogen *L. biflexa*, which lacks the *sph2* gene.

As shown in Fig. 5A, *gfp* expression from the fusions in the Lai strain reflected the expression observed with native *sph2* in the Lai and Pomona strains in a previous study (10). GFP levels from the *sph2p*_Lai_*-gfp* fusion in the Lai strain incubated in EMJH were near background levels. Expression from the *sph2p*_Pom_*-gfp* fusion in was much higher than from the *sph2p*_Lai_*-gfp* fusion (Fig. 5A). The same two fusions were also examined in the Pomona strain. In contrast to its activity in the Lai strain, the *sph2p*_Lai_*-gfp* fusion produced large amounts of GFP in the Pomona strain, although not as much as the *sph2p*_Pom_*-gfp* fusion (Fig. 5B). These observations suggest that the activity of the (Lai) *sph2* promoter was being suppressed in the Lai strain. Neither *sph2* promoter fusion produced GFP above background levels in *L. biflexa* (Fig. 5C). GFP activity was detected with the control *lipL41p-gfp* fusion in all three strains (Fig. 5).

**FIG 5.**
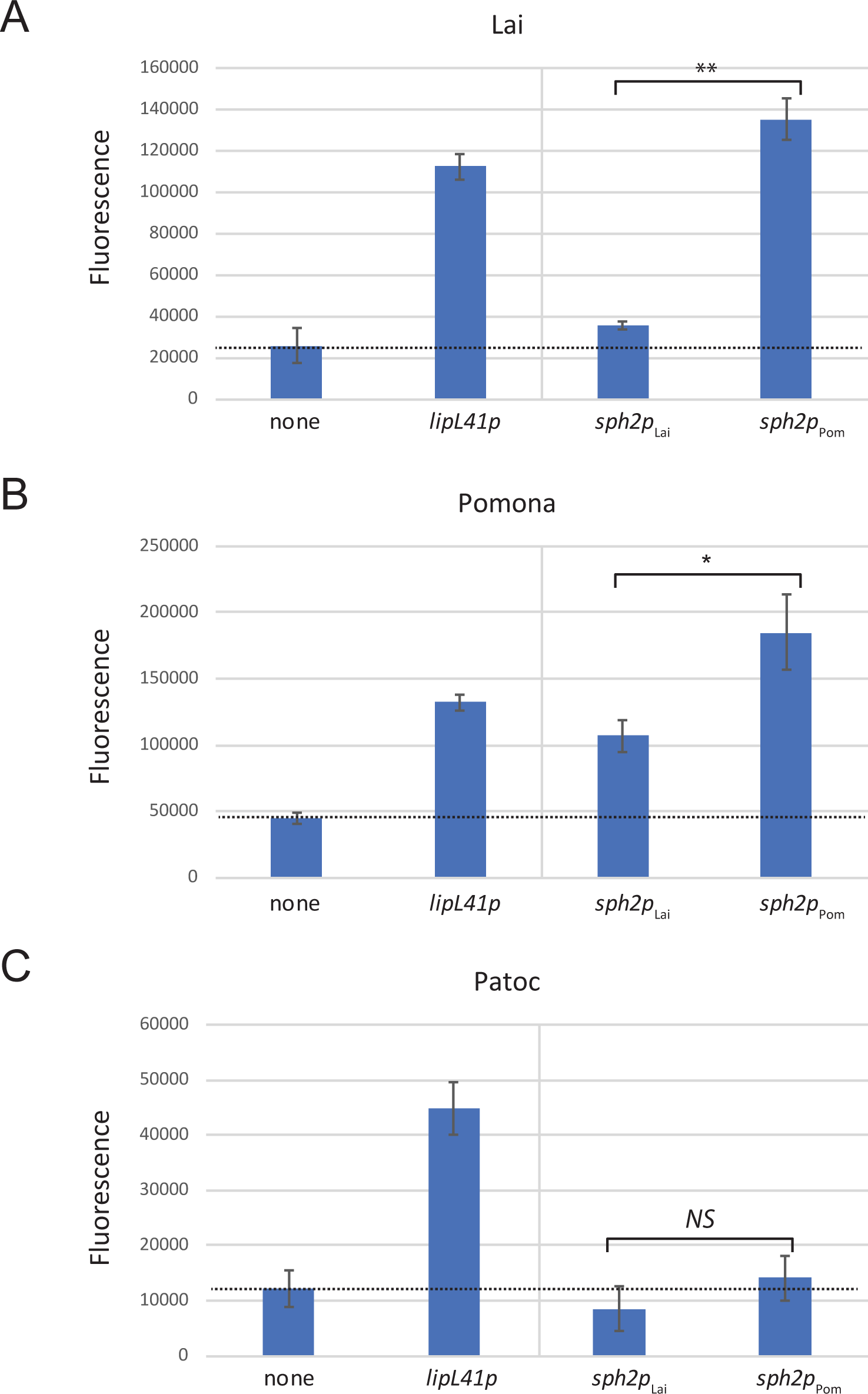
GFP production from *sph2p-gfp* fusions in *L. interrogans* and *L. biflexa*. Fusion plasmids were introduced into *L. interrogans* Lai strain 56601 (Lai), *L. interrogans* Pomona strain LC82-25 (Pomona), and *L. biflexa* strain Patoc I (Patoc) and incubated in EMJH with spectinomycin. The dashed line indicates the mean background reading obtained with transconjugants carrying the promoterless *gfp* fusion vector pRAT724 (none). *P* values were calculated from Welch’s t-test: *, *P* < 0.05; **, *P* < 0.01, *NS*, not significant.

We further examined the activities of the promoters in the Lai strain under different conditions of growth. To examine fusion expression at different cell densities, cultures of the four Lai fusion strains were started at a low cell density. When the density reached 1 x 10^7^/ml, GFP production from the fusion strains was examined daily until early stationary phase was reached. For all fusions, GFP levels rose as the cultures reached stationary phase, although the rise was slightly greater for the *sph2* promoter from the Pomona strain and the *lipL41* promoter (Fig. 6). Notably, GFP from the Lai *sph2* promoter fusion never rose above background levels (Fig. 6).

**FIG 6.**
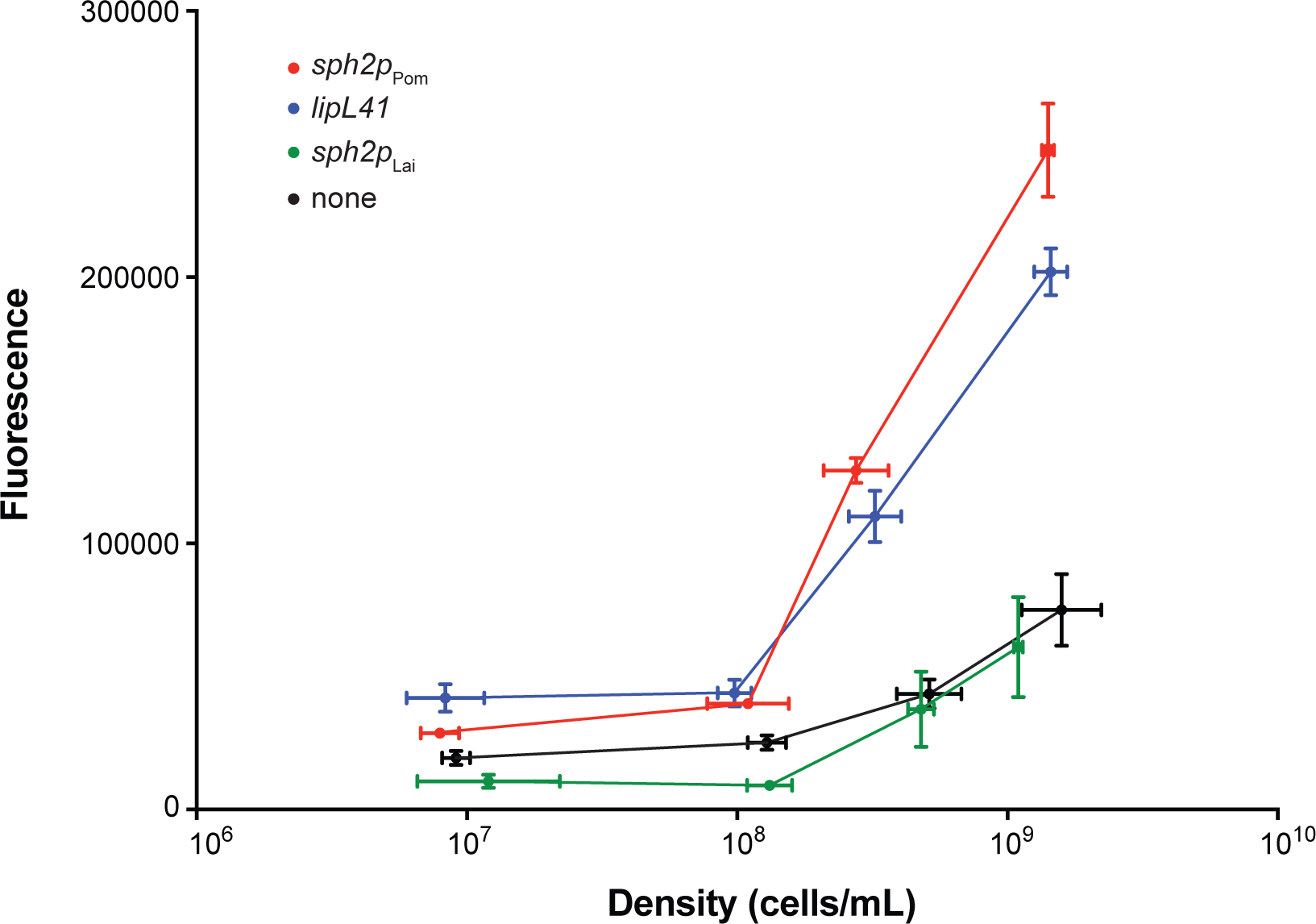
GFP production from *sph2p-gfp* fusions in *L. interrogans* at different cell densities. Cultures of tranconjugants harboring the fusion plasmids were sampled daily starting from a cell density of 1 x 10^7^/ml. All samples were assayed for fluorescence.

We next determined whether the increase in native Sph2 levels observed when *L. interrogans* is shifted from EMJH to EMJH with sodium chloride added to physiologic osmolarity (Narayanavari 2015) is also observed with the *sph2* fusions. Fusion expression was higher when the transconjugants harboring the *sph2p-gfp* constructs were grown in EMJH with sodium chloride added to physiologic osmolarity (Fig. 7). The *sph2p*_Pom_*-gfp* fusion generated high levels of GFP in the Lai strain, and expression was even higher at physiologic osmolarity (Fig. 7). As expected, expression from the control *lipL41p-gfp* fusion was not affected by the sodium chloride supplement (Fig. 7).

**FIG 7.**
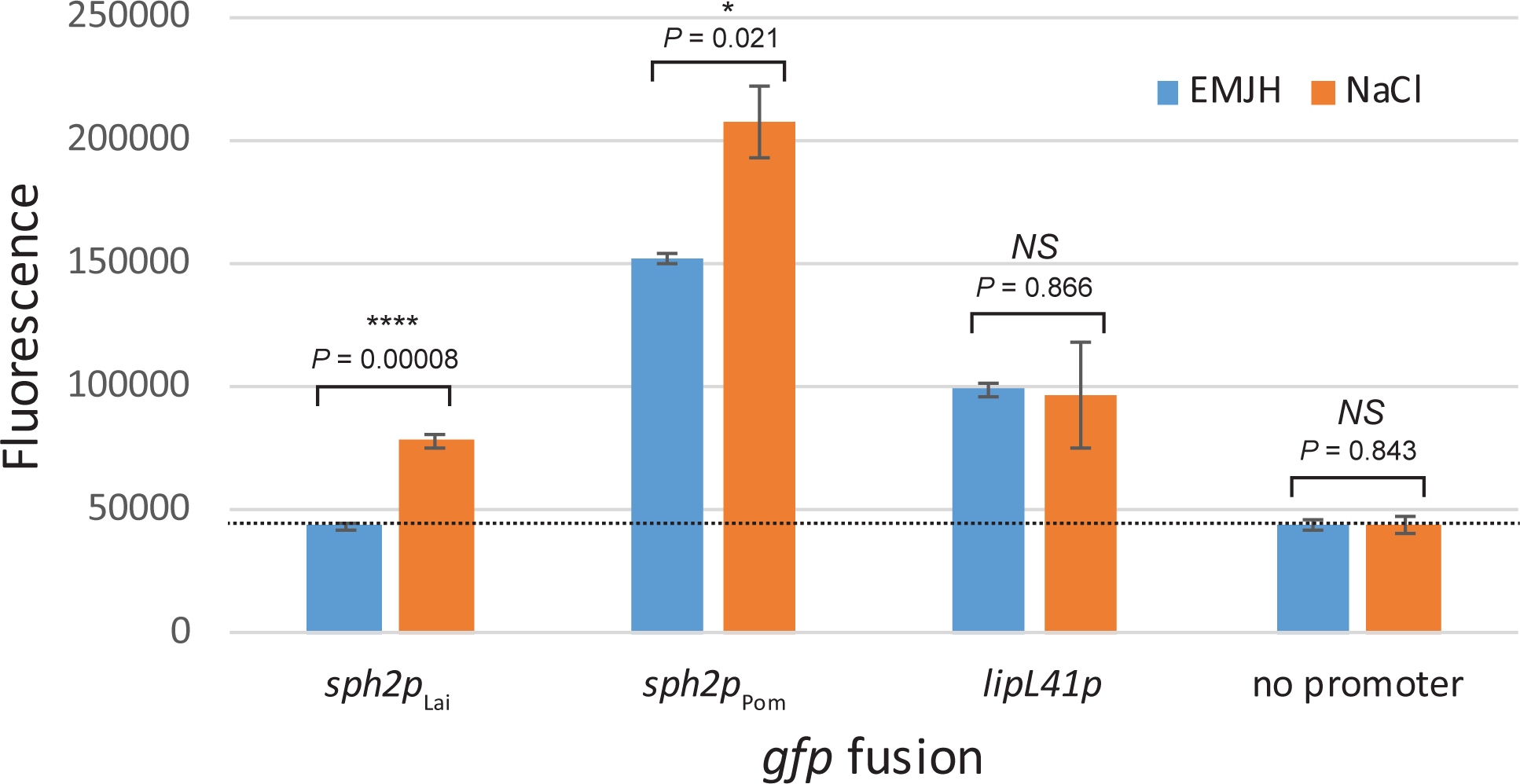
Regulation of *sph2p-gfp* fusion expression by sodium chloride. Fusion plasmids were introduced into *L. interrogans* Lai strain 56601. Flourescence was measured from transconjugants maintained in EMJH or EMJH with 120 mM sodium chloride for 4 hours. Means ± SD are plotted (n = 3). *P* values were calculated from the Welch’s t-test comparing fluorescence with and without 120 mM sodium chloride added to EMJH (panel A) or with and without the IS-like element (panel B): *, *P* < 0.05; ****, *P* < 0.0001; *NS*, not significant.

To determine whether the IS-like element in the promoter region of *sph2* of the Pomona strain is responsible for its high basal level of expression, the IS-like element (Fig. 8A) was precisely deleted from the *sph2* promoter region of Pomona, and the promoter variant was fused to *gfp*. Surprisingly, the GFP assay showed that removal of the IS-like element had no significant effect on expression of the *sph2* promoter fusion when examined in the Lai strain incubated in EMJH medium and in EMJH supplemented with sodium chloride (Fig. 8B).

**FIG 8.**
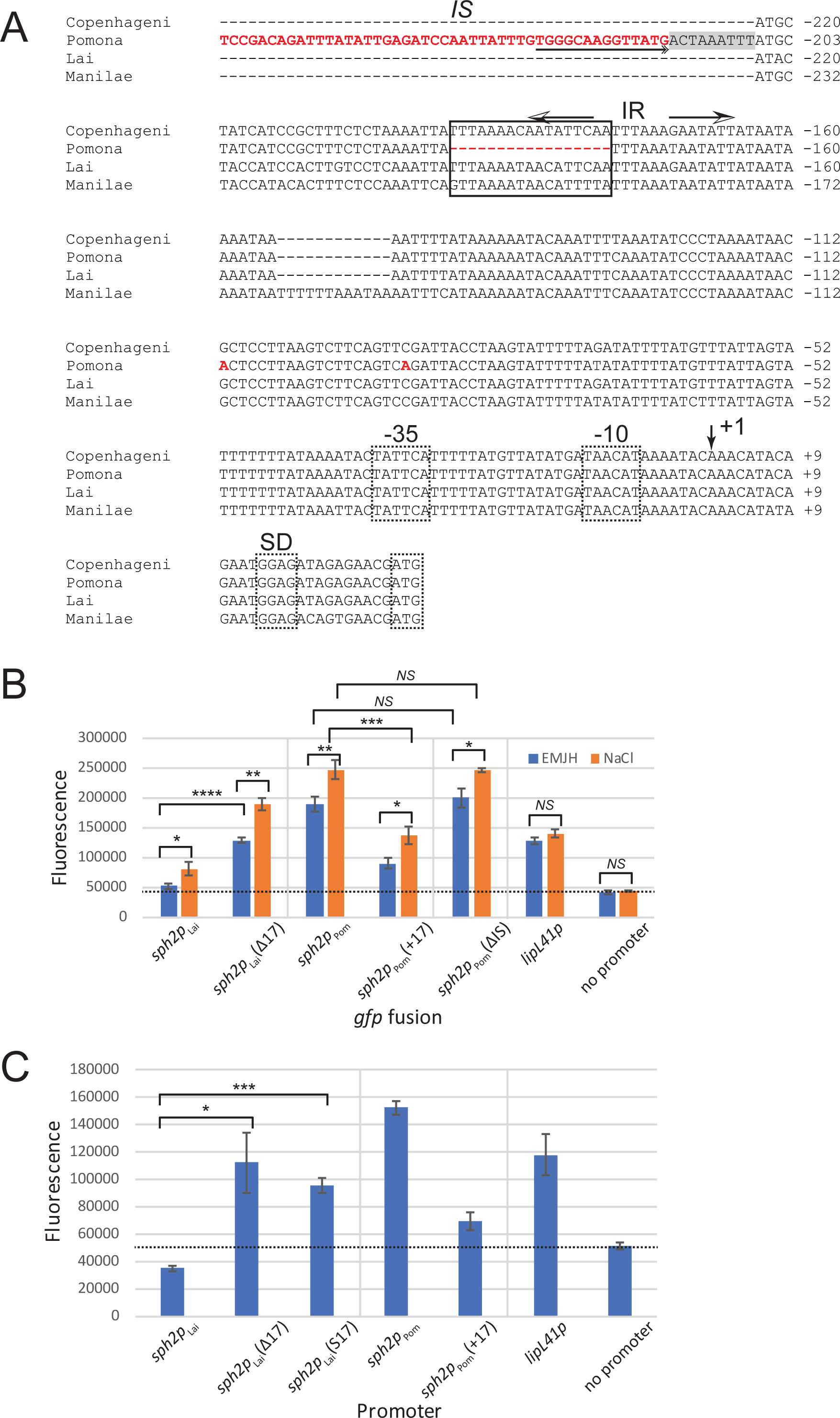
Effect of mutations introduced upstream of the *sph2* promoter. A. Multisequence alignment. Sequences upstream of *sph2* from *L. interrogans* strains Fiocruz L1-130 (Copenhageni), LC82-25 (Pomona), 56601 (Lai), and L495 (Manilae) were aligned with Kalign. The aligned sequences extend from the 5’ end of the promoter fragments cloned into pRAT724 to the start codon of *sph2*. Most of the IS-like sequences (*IS*) and sequences further upstream were omitted from the figure for brevity. Sequences unique to the Pomona LC82-25 are shown in red. An outside end of the IS-like element containing an inverted repeat is underlined with a double arrowhead at the end. The duplicated host sequence adjacent to the insertion site of the IS-like element is shaded gray. The key 17-nucleotide sequence is boxed, and the imperfect inverted repeats (IR) overlapping with the key sequence are marked with arrows. The proposed transcription start site is indicated by an arrow at position +1. The *E. coli*-like -10 and -35 promoter sequences, Shine-Dalgarno sequence (SD), and start codons are demarked with dashed boxes. B and C. Fusion expression. Fusion plasmids were introduced into *L. interrogans* serovar Lai strain 56601. Transconjugants were incubated in EMJH or EMJH with 120 mM sodium chloride for 4 hours (panel B) or left in EMJH (panel C). The dashed line indicates the mean background reading obtained with *L. interrogans* carrying the empty *gfp* fusion vector pRAT724. Means ± SD are plotted (n = 3). *P* values were calculated with the Welch’s t-test: *, *P* < 0.05; **, *P* < 0.01; ***, *P* < 0.001; ****, *P* < 0.0001.

To search for sequences that could account for the differences of basal *sph2* expression between the Pomona LC82-25 strain and non-Pomona strains, we aligned the sequences upstream of the *sph2* coding regions from the Pomona LC82-25, Lai 56601, Copenhageni Fiocruz L1-130, and Manilae L495 strains. The 5’ end of the aligned sequences correspond to the 5’ end of the *sph2* promoter fragments cloned into pRAT724. As expected, the multisequence alignment revealed a 321 bp IS-like sequence in the Pomona strain located from position -216 through -536 relative to the major 5’ end of the *sph2* transcript of the Pomona strain (Fig. 8A) (13). Sequence elements characteristic of transposable elements were present: a 17 bp inverted repeat at the ends of the element and a 9 bp direct repeat of host sequences immediately adjacent to the inverted repeat (Fig. 8A).

An additional unique feature of the Pomona sequence is its lack of a 17-base pair segment between positions -178 and -179 relative to the major 5’ end of its *sph2* transcript (Fig. 8A). To determine whether this 17 bp sequence was responsible for the low basal *sph2* expression in the Lai strain, the 17 bp sequence was removed from the upstream region of the Lai *sph2* promoter in the *gfp* fusion. The deletion mutation restored 58% (Fig. 8B), 63%, and 81% of the expression observed with the Pomona *sph2p-gfp* fusion over three experiments. The converse experiment was also performed by inserting the 17 bp segment upstream of the Pomona *sph2* promoter in the fusion. The insertion reduced fusion expression by 68% (Fig. 8B), 70%, and 72%. Expression from all *sph2* fusions was elevated in the presence of physiologic osmolarity, indicating that the 17 bp sequence is not a major contributor to osmotic induction of *sph2* expression.

To rule out the possibility that removing the 17 bp sequence altered spacing between transcriptional control elements, an additional *sph2* promoter variant was constructed by scrambling the 17 bp sequence. The scrambled sequence restored 44% of the expression observed with the Pomona fusion (Fig. 8C), indicating that at least a portion of the 17 bp sequence plays a direct role in limiting transcription from the *sph2* promoter.

## DISCUSSION

Differential expression of the *sph2* sphingomyelinase gene from a Pomona and a non-Pomona strain of *L. interrogans* was examined with a new *gfp* fusion plasmid. This *gfp* fusion plasmid enabled identification of a segment of the Lai strain’s *sph2* promoter that is primarily responsible for its low levels of *sph2* expression. The Copenhageni and Manilae strains, which also express low basal levels of *sph2*, also possess the 17 bp sequence. The three Pomona strains, which express high levels of Sph2, lack the 17 bp segment. Interestingly, the 17 bp sequence contains one arm of an inverted repeat (Fig. 6A), which may be a binding site for a *trans*-acting factor responsible for the low basal expression of *sph2*. In contrast, an upstream IS-like sequence hypothesized to increase *sph2* expression did not contribute to basal expression from the Pomona *sph2* promoter when examined in the Lai strain.

It is anticipated that this *gfp*-reporter strategy can be used to study gene expression in other strains and species of *Leptospira*. The parent of pRAT724, pMaORI, stably replicates in at least five other strains of *L. interrogans*, *L. mayottensis*, *L. licerasiae*, *L. fainei*, and *L. biflexa* (24). pMaORI carrying the *fliM* gene also restored the virulence of a spontaneous motility mutant of *L. interrogans*, indicating that pMaORI is stable *in vivo* in the absence of antibiotic selection (36). Nevertheless, future efforts to examine expression of promoter fusions to *gfp in vivo* will require additional experiments to verify stability of the fusion plasmid in the absence of antibiotics. One limitation of the new fusion plasmid is that variability and irreproducibility in the GFP levels were observed at high levels of expression of the *sph2* promoter fusions in the Pomona strain. For this reason, although the IS-like element did not contribute to *sph2* expression in the Lai strain, we cannot firmly exclude the possibility that the IS-like element contributes to basal *sph2* expression in the Pomona strain. Nevertheless, development of this *gfp* fusion plasmid will facilitate identification of *cis*-acting sequences critical for expression of *L. interrogans* genes. The recent publication of the genome-wide transcription start sites for *L. interrogans* will aid selection of the correct junction for promoter fusions to *gfp* (37). A transcription start site was not detected for *sph2* in the study, consistent with the limited expression of *sph2* in non-Pomona strains under standard culture conditions.

Elucidation of the mechanism for differential *sph2* gene expression among *L. interrogans* strains raises the interesting question of whether the 17-bp promoter deletion provides a selective benefit to these strains. Increased *sph2* expression appears to be unique among Pomona strains (10, 11,). In contrast, *sph2* is expressed poorly in strains from serovars Copenhageni, Lai, and Manilae, whose reservoir hosts are small rodents (38-40). The association of serovar Pomona infection with sheep and pigs (9) suggests that high levels of *sph2* expression confer an adaptive benefit to the Pomona serovar for infection of specific maintenance hosts. Additional studies comparing infection in different hosts with leptospiral strains with and without the 17-bp deletion would be helpful in understanding the role of Sph2, assessing whether there is a benefit of increased *sph2* expression in some hosts, and determining whether that benefit is host specific.

The *sph2* fusion constructs described here may be helpful in identifying the *cis*-acting site(s) involved in osmotic regulation of *sph2*. In a previous study, addition of 120 mM sodium chloride to cultures increased *sph2* transcript levels by over 100-fold in both serovar Manilae and serovar Pomona (10). The fact that these increases were similar in both low- and high-Sph2 producers suggests that the 17-bp *cis-*acting element is not involved in osmotic regulation of transcription. A previous study with the 600 bp immediately upstream of the *sph2* coding region from the Fiocruz L1-130 strain of *L. interrogans* fused to *gfp* demonstrated osmotic induction of the fusion in *L. biflexa* (18). We observed osmotic induction with a fragment extending from 24 to 325 nucleotides upstream of the *sph2* coding region fused to *gfp*. Therefore, the 302 bp region serves as a starting point to locate the regulatory target. The observation that osmotic induction occurs in the nonpathogen *L. biflexa* suggests that a *trans*-acting determinant of *sph2* osmotic regulation is conserved between at least some pathogenic and nonpathogenic *Leptospira* species.

Little is known about how *cis*- and *trans*-acting factors establish the level of gene expression in *Leptospira*. Several *cis*-acting sequences affecting expression of *Leptospira* genes involving adaptation to different environmental stresses have been identified. The *L. interrogans* genome encodes two *lexA* genes (41). DNase I footprinting and gel-shift assays with the *recA* upstream sequence revealed an SOS box bound by LexA1 (42). Additional gel-shift assays demonstrated that similar sequences upstream of other SOS-induced *L. interrogans* genes bound to LexA1 (41). LexA2 bound only to its own promoter (41). In another study, a high-affinity binding site for the phosphorylated form of the *L. biflexa* HemR response regulator, which regulates heme metabolism, was enriched from a highly-randomized oligonucleotide library by several rounds of affinity selection with the phosphorylated protein (43). In a recent study, the *L. interrogans* enhancer binding protein EBP and the alternative sigma factor RpoN was shown to bind to various promoters by gel shift analysis (44). For *L. interrogans* genes such as *sph2* for which the *trans*-acting regulatory factor remains to be identified, the pRAT724 *gfp* reporter plasmid will facilitate identification of additional *cis*-acting sequences affecting gene expression.

## ACKNOWLEDGEMENTS

We thank William G. Miller for providing pGreenTIR and Mathieu Picardeau for contributing pMaORI. We also thank Rich Zuerner for supplying the serovar Pomona strains of *L. interrogans*. We thank Suneel Narayanavari for technical assistance and David Beenhouwer for use of the fluorescence microplate reader.

## Funding

The study was funded by Veterans Affairs Merit Awards (to J.M. and D.A.H.) and Public Health Service grant R21 AI128560-01A1 from the National Institute of Allergy and Infectious Diseases (to D.A.H).

## Author contributions

J.M. conceived the study, contributed materials, designed and performed the experiments, analyzed the data, and wrote the paper. D.A.H. helped analyze the data and write the paper.

